# Do higher-order moments improve inference of population dynamics?

**DOI:** 10.64898/2026.09.28.754680

**Authors:** Chandan Relekar, Amanda de Azevedo-Lopes, Michael Sieber, Arne Traulsen

## Abstract

Deciphering the interactions among host-associated microbial communities, such as the gut microbiome, is an important, but challenging task — there is constant flux of microbes and frequent environmental fluctuations driven by external factors and host immune response. At the same time, data collection is inherently constrained, as non-invasive sampling methods only capture snapshots of the system, and the number of available sampling points is often small. In such non-ideal cases, Bayesian parameter inference allows us not only to estimate model parameters from limited and noisy data, but also to quantify the uncertainty in those estimates. Studies using Bayesian techniques typically focus on fitting the mean dynamics to averaged data alone, neglecting the information provided by higher-order moments. In this study, we fit stochastic models of population dynamics to simulated microbiome data, and investigate whether the additional fitting of second-order moments, that is, fitting the variance and covariance observed across replicates to those expected from demographic stochasticity in the underlying model, can improve parameter inference by incorporating information that would otherwise be discarded. For simulated datasets, we observe a substantial improvement in parameter inference. However, we find that both the manner in which the first-order moments and second-order moments are combined in the distance function and the stopping criterion used during the optimization procedure substantially influence the quality of parameter inference.

**Author summary:** Fitting mathematical models of population dynamics to microbial time-series data allows us to estimate the ecological processes and interactions taking place in the microbiome. Repeated experiments of microbial systems yield replicates which slightly differ from each other. Some of this variability arises due to the fact that births and deaths occur at random. Most prior work focuses on fitting a deterministic mathematical model to the average across replicates. We use a stochastic model to fit the variability to the observed variability across replicates. Using a simulation-driven approach, we study the conditions under which our approach allows us to infer a larger fraction of ecological parameters correctly. We observe a substantial improvement in parameter inference. Lastly, our Bayesian approach not only allows us to incorporate prior information about the system, but also provides a distribution of parameters which conveys some idea of the uncertainty of the estimates.

## 1 Introduction

Host-associated microbial communities, such as the gut microbiome, have gained a lot of attention in recent years due to their influence on host health and disease [1–4]. There is a great interest in inferring the underlying ecological interactions and processes within these communities (refer to Ref. [5] for a review of available tools), which could give us the ability to manipulate them to a more desirable state for example via antibiotics, diet, or fecal-microbiota transplantations. One of the approaches to infer these interactions and processes is to fit a mathematical model to gut-microbiome data and estimate the model parameters that best explain the observed time-series. There has been extensive literature that fit versions of the deterministic Lotka-Volterra model to simulated and empirical data using regression methods, such as least squares or maximum likelihood estimation, which yield point-estimates of the interactions [6–8]. While this can sometimes lead to accurate time-series predictions, the estimation of the underlying parameters can be erroneous [6]. Due to the complex dynamics and the associated stochastic effects, accurate model predictions are fundamentally challenging.

Zapién-Campos and colleagues have explored the use of stochastic models to infer parameters from microbiome time-series data, while also extracting additional information from the data beyond the mean dynamics, by deriving and fitting higher-order moments of the stochastic model to the data [9]. Here, we build on this framework and systematically explore the extent to which, and the conditions under which, the addition of higher-order moments improves parameter inference.

Using a Bayesian approach, we obtain posterior probability distributions over parameter values, providing probabilistic estimates of what the parameters are likely to be. Bayesian approaches have been used to fit models to data [10–12] and some studies show that:

1. Stochastic models are better identifiable [13, 14] - both structurally (in the face of infinite, noiseless data) and practically (i.e. with real-world constraints).
2. Fitting higher-order statistical moments (e.g., fitting the variances and covariances observed in data to what is expected from demographic stochasticity in the underlying model, in addition to fitting the average behavior across replicates) leads to better parameter inference [15]. Besides, higher-order moments are better approximations of stochastic systems at the mesoscopic scale [16].
3. Bayesian methods have been found to perform better at parameter inference with irregularly and sparsely sampled time-series data, which is often the case with microbiome datasets [10].

Directly using a stochastic model to fit to data is computationally expensive due to the sheer number of stochastic trajectories that need to be simulated during the inference procedure. Thus, fitting statistical moments of an underlying stochastic model to data has an additional benefit of decreasing the computational cost. We combine the approaches listed above and determine if and to what extent they improve our ability to infer parameters. We focus on characterizing the conditions under which incorporating higher-order moments improves parameter identifiability and inference accuracy. Namely, we characterize the impact of the number of microbial types on the inference quality with absolute and relative abundances, study the impact of how information from first-order moments and second-order moments are combined in the distance function during the fitting procedure, and the effect of different stopping criteria during the optimization procedure. For simulated datasets of absolute abundances, a larger fraction of parameters is correctly inferred

when we use both first- and second-order moments instead of first-order moments only. Moreover, parameters associated with more abundant types tend to be inferred more accurately than parameters associated with less abundant types. Inference is more difficult when working with relative abundances, indicated by a lower fraction of parameters that is accurately inferred – regardless of the inclusion of second-order moments.

## 2 Methods

We build on the computational workflow from Ref. [9], where we start with microscopic transition rates describing changes in the microbiome composition of one host. From these rates, we can describe how the probability of a microbiome composition in an ensemble of hosts changes over time, which is described by the master equation [17, 18]. The master equation allows us to derive equations for the statistical moments of the microbiome population dynamics, which we can then use to fit to data and infer the parameters of the simulated datasets. A schematic of the workflow is shown in Figure 1. We start with the formulation of the transition rates, the statistical moments for the logistic and Lotka-Volterra models derived in Ref. [9], and the Bayesian inference procedure used to fit the models to the data.

**Figure 1:**
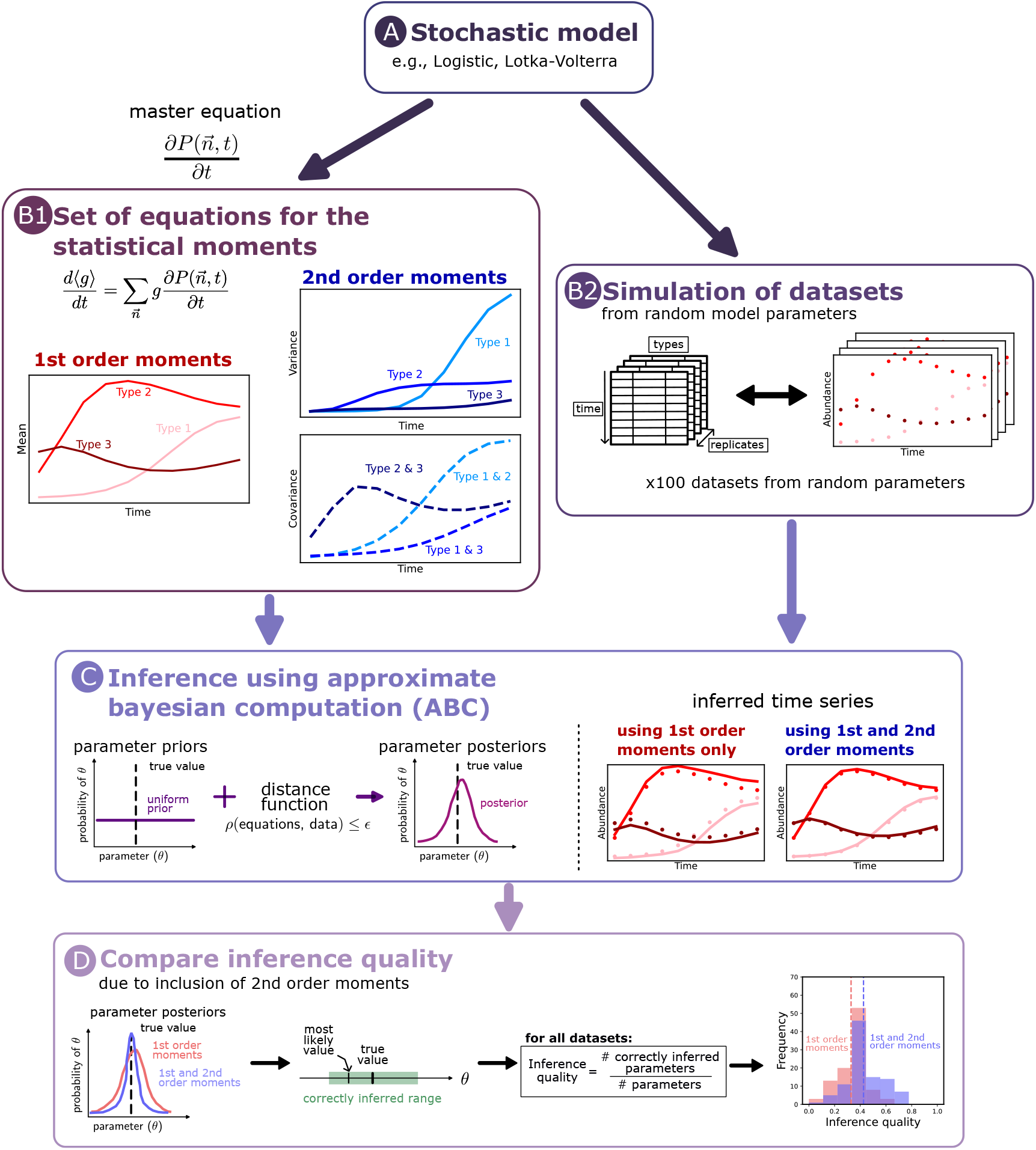
Schematic of the workflow. Starting from the stochastic model (A), the corresponding equations for the statistical moments are derived from the master equation (B1). Here, we consider both first-order (mean) and second-order (variance and covariance) moments. Using the stochastic model, we generate datasets containing absolute or relative abundances by simulating four replicates of the system using randomly sampled model parameters (B2). Using Approximate Bayesian Computation (C), the derived statistical moments are fit to the generated time-series data, and the model parameters are extracted. We then systematically evaluate to what extent the inclusion of higher-order moments improves the accuracy of parameter inference, across a range of number of types and data types (D).

### 2.1 Logistic model

In the logistic model with *L* types, each type *k* (*k* = 1, …, *L*) has a growth rate *f*_*k*_, a death rate *ϕ*_*k*_ and an immigration rate *m*_*k*_. We assume a shared carrying capacity *N* above which no growth takes place. We assume that only a single microbe can divide or die at a given point of time. Thus, the number of microbes of type *k* increases at rate (*f*_*k*_*n*_*k*_ + *m*_*k*_) (1 − ∑ _*j*_ *n*_*j*_*/N*) and decreases at rate *ϕ*_*k*_*n*_*k*_*/N*. In a notation that we will also use later, the transition rates are

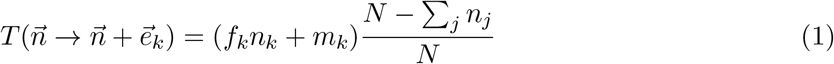

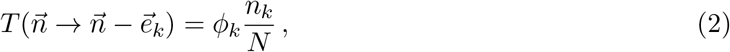

where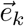 is a vector with one in the *k*^*th*^ entry and zeros elsewhere.

For each model with *L* types, we randomly sample 100 parameter sets. For each parameter set, we simulate synthetic time-series data using the Gillespie algorithm [19] and the transition rates in Eqs. (1) and (2) to generate four independent replicates. The generation of synthetic time-series is described in Appendix E.

Using the above expressions, we can write the master equation [18]

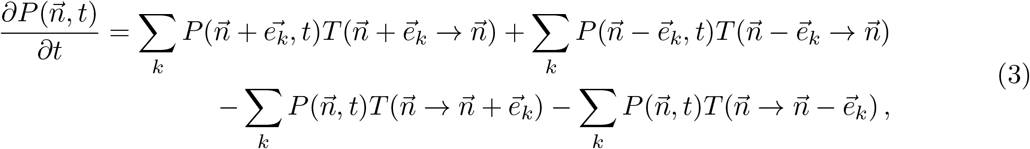

which is used to derive an analytical expression that describes the dynamics of the average of a given dynamical variable *g*,

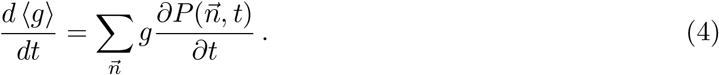

With *g* = *n*_*k*_, we obtain [9]

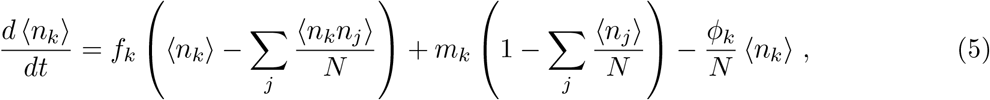

which are the *L* first-order statistical moments. Similarly, there are *L*^2^ equations for the second-order statistical moments. They can be derived by setting 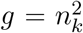 and *g* = *n*_*k*_*n*_*l*_, giving rise to *L* second-order moments and *L*^2^ −*L* co-moments. The derived expressions for the statistical moments can be found in Appendix A.1.

### 2.2 Lotka-Volterra model

In the Lotka-Volterra model, each type *k* has a maximum growth rate *f*_*k*_. The interaction matrix *I*, is split into positive interactions *A* and negative interactions *B. A* and *B* have positive entries only. In contrast to the logistic model, where the interactions between types are neutral, the Lotka-Volterra model includes directional interactions between every pair of types. The transition rates for the Lotka-Volterra model with *L* types are

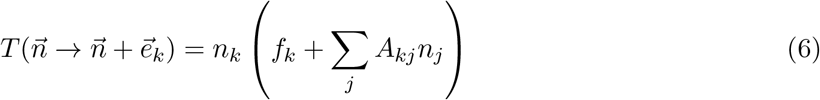

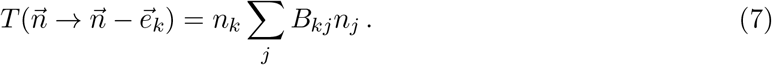

Repeating the same procedure as before, we obtain equations for *L* first-order moments and *L*^2^ second-order moments. The derived expressions can be found in Appendix A.2.

### 2.3 Moment closure

The equation for the first-order moments ⟨*n*_*k*_⟩ (for example, Eq. (5)) contains the second-order terms, ⟨*n*_*k*_*n*_*l*_⟩. In general, the equations of the *n*-th order moments contain terms of the (*n* + 1)-th order. This makes simulation and fitting of the system impossible. One approach is to ‘close’ the system at a certain order [20]. An *n*-th order moment-closure approximation [21] involves breaking down terms of (*n* + 1) order into the lower order [22, 23]. For example, the first-order moment-closure approximation applied on equation (5) involves using the approximation - ⟨*n*_*k*_*n*_*l*_⟩ ≈ ⟨*n*_*k*_⟩ ⟨*n*_*l*_⟩. The second-order moment-closure approximation applied on the entire system requires using the approximations 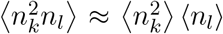 and ⟨*n*_*k*_*n*_*l*_*n*_*j*_⟩ ≈ ⟨*n*_*k*_*n*_*l*_⟩ ⟨*n*_*j*_⟩ on equations (S5) and (S6) while leaving equation (5) untouched, since it does not feature any terms above second-order. Without moment-closure approximations, one would have to rely on exhaustive stochastic simulations in the parameter space during the optimization procedure, which turns out to be computationally demanding. Moment-closure approximations can substitute stochastic simulations under certain conditions [16].

### 2.4 Distance functions

Let 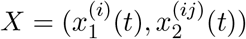 and 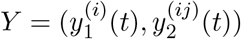 be two time-series, where subscript 1 denotes the first-order moments, and subscript 2 denotes the second-order moments. The sampling times are *T* = *{t*_0_, *t*_1_, …, *t*_*m−*1_*}*. The distance function *ρ*(*X, Y*) returns, at each time point *t*, a number that describes the distance between the two time-series *X* and *Y*. These values are then averaged over all *t*. There are different ways of combining information from the first-order moments and second-order moments. The order of magnitude of the second-order moments is twice that of the first-order moments in the case of absolute abundances, and half that in the case of relative abundances. Hence, in order to enable a meaningful comparison between the cases with and without inclusion of second-order moments, the second-order moments are rescaled before combining them with the first-order moments in the distance function. Table 1 shows different ways of combining the first-order moments and second-order moments in the distance function *ρ*(*X, Y*).

**Table 1:**
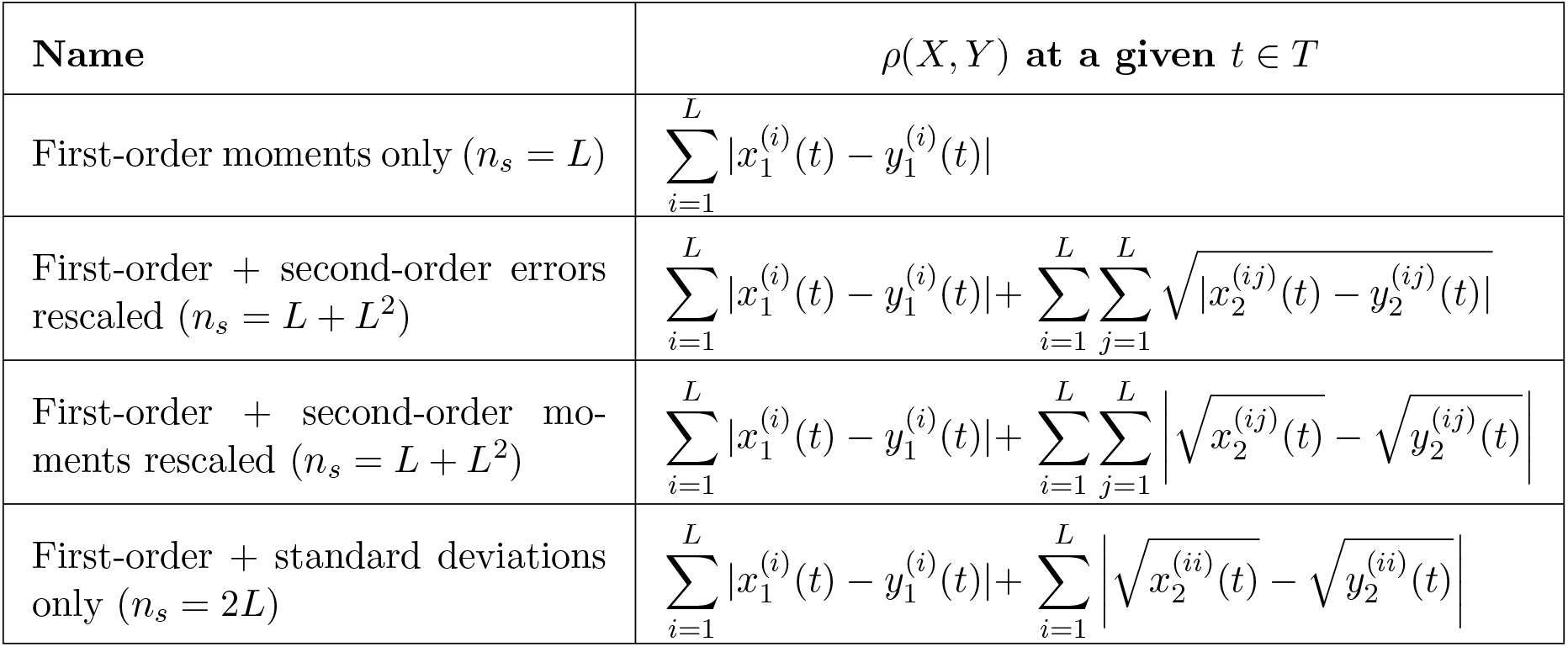
Summary of the used distance functions, *ρ*(*X, Y*). Dissimilarity between the first-order moments and second-order moments can be combined in different ways when computing the distance function, while making sure that the second-order moments do not dominate the overall distance function. *n*_*s*_ is the total number of terms inside the summations.

### 2.5 Parameter inference

We implemented our parameter inference workflow using Approximate Bayesian Computation (ABC) with Sequential Monte Carlo (SMC) [24]. The workflow requires:

1. Data *d* that has to be fitted to the model. In our case, the data *d* is a time-series that is either synthetically generated or obtained experimentally.
2. A model function *H* that takes in the model parameters *θ* and returns an output *H*(*θ*) that is plugged into the distance function. In our case, the model functions are the ODEs of the logistic and Lotka-Volterra models that generate time-series using the given model parameters *θ*.
3. A distance function *ρ*(·, ·) is used to compare (and therefore minimize) the distance between the model output and the data.
4. Priors for the parameters that have to be estimated, which represent any previous information about the distribution of parameter values. We have used uniform (uninformative) priors throughout, which have minimal influence on inference beyond conveying the range in which the parameters likely exist. ABC iteratively generates posterior distributions for the parameters which combines the numerically estimated likelihood and the priors. The used priors are listed in Appendix C.

This approach iteratively generates posteriors from the previous priors based on what parameter distributions are more likely to minimize *ρ*(*H*(*θ*), *d*), until the distance falls below a set threshold *ϵ*. Our formulation of the distance function allows us to set *ϵ* in an intuitive manner, such that it describes the tolerated error per time-series per time-point, between the data and the final posterior distribution. We implemented this workflow using tools from the Python package pyABC [25]. Additionally, ABC involves a careful calibration of several hyper-parameters, which are discussed in Appendix D.

### 2.6 Accuracy metric

An accuracy metric is required for gauging the performance of our parameter inference framework on synthetic data generated using known parameter values. The final output of ABC are posterior distributions for every parameter. The extent to which the posterior has collapsed is a measure of precision of the prediction. We treat the most-likely value as the prediction of the framework. If the predicted value is within 10% of the true value we deem that the parameter has been predicted correctly. The inference quality is the fraction of parameters that have been predicted correctly averaged across all 100 randomly generated time-series.

## 3 Results

### 3.1 Evaluation on synthetic datasets

We evaluated the performance of our parameter inference framework on synthetic data generated using known parameter values. For both the logistic and Lotka-Volterra models, and for each number of types *L*, we generated 100 randomized datasets and compared the inference quality of fitting only first-order moments to fitting both first- and second-order moments (Figure 2). With absolute abundances, we find that fitting both first- and second-order moments on average leads to a higher fraction of parameters that are inferred correctly compared to using just first-order moments. This is seen across both the logistic and Lotka-Volterra models. However, when working with relative abundances, the inference quality does not increase in the same way (Figure S1 and Appendix B). In both models, the inference quality decreases as the number of types increases (Figure 2), even under our idealized conditions where the synthetic data is free of measurement noise and features only demographic stochasticity.

**Figure 2:**
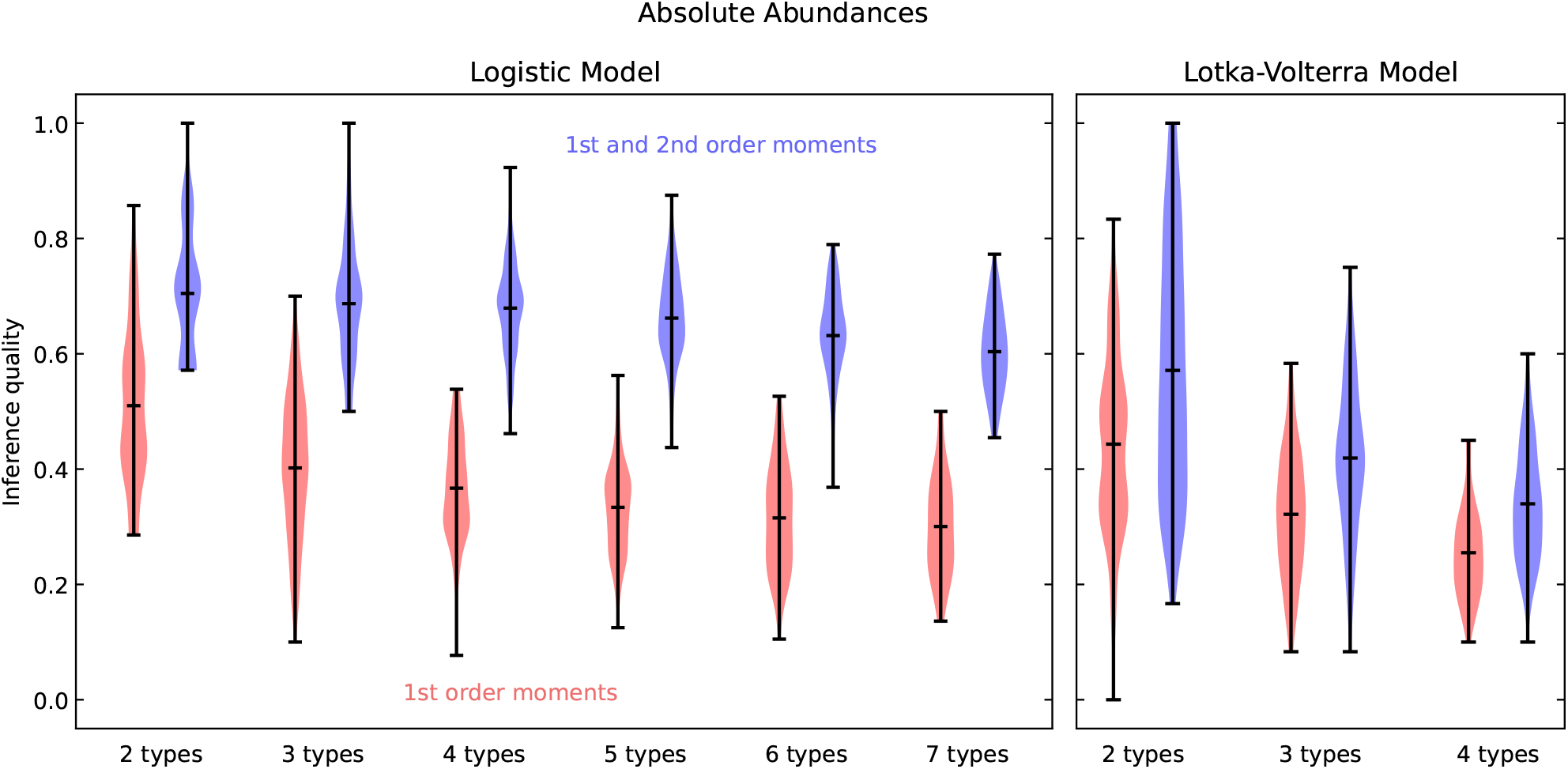
Estimating the inference quality on synthetic data with absolute abundances. We compare the inference quality of fitting only first-order moments to the inference quality of fitting both first- and second-order moments, across 100 randomized datasets, for the logistic and the Lotka-Volterra models and different numbers of types, *L*. Across the board, we observe an increase in inference quality due to incorporation of second-order moments. In both models, the inference quality decreases as the number of types increases. For the first-order case, the ‘first-order moments only’ distance function is used. For the first- and second-order case, the ‘first-order and second-order errors rescaled’ distance function is used. Both distance functions are listed in Table 1.

We also study how the inference quality varies as a function of the error threshold utilized during inference. Figure S4 showcases this for the case of the logistic model with five types. The inference quality obtained by fitting only first-order moments to a very low error threshold is comparable to the inference quality achieved when additionally fitting second-order moments to a higher error threshold. However, fitting to such low error thresholds is only feasible with synthetically generated datasets, whereas empirical datasets not only contain noise but may also exhibit substantial deviation from theoretical models.

In Figure 3, we show the posterior distributions obtained by fitting the Lotka-Volterra model with 3 types and absolute abundances to a representative time-series. Not only are the inferred parameters more accurate with the incorporation of second-order moments (as shown in Figure 2), but the inference also becomes more precise, as indicated by the narrower posterior distributions in Figure 3. Figure S6 contains the posterior distributions for the logistic model, analogous to the results in Figure 3. The growth rates, immigration rates, and carrying capacity are more accurate and the distribution narrows with the incorporation of second-order moments. However, the death rates are not accurately identified (Figure S5). We show the original and predicted time-series corresponding to the posterior distributions in Figure 3 and Figure S6 in Figure S7.

**Figure 3:**
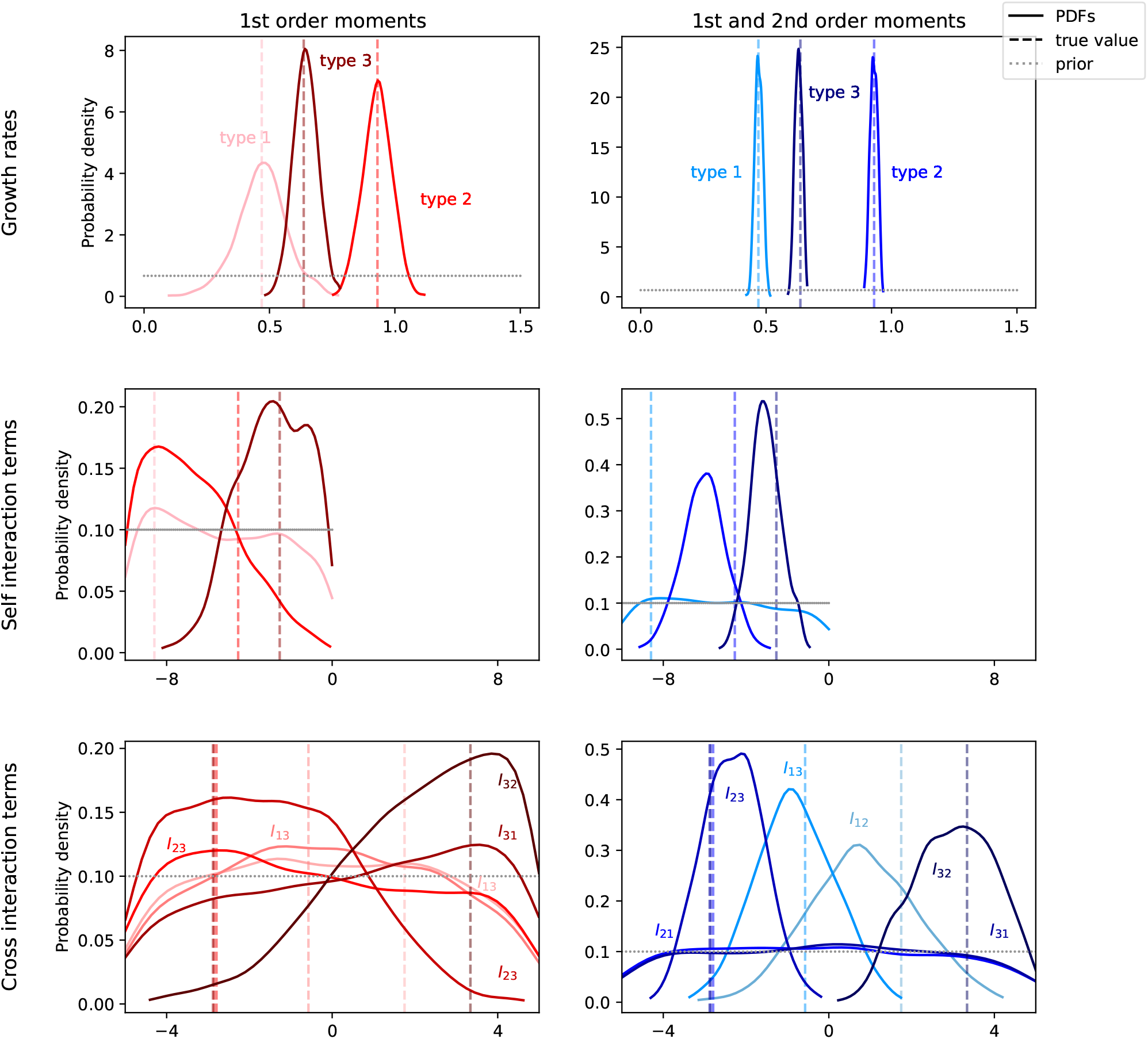
Comparison of the posterior distributions with and without the inclusion of second-order moments for the Lotka-Volterra model with three types and absolute abundances. All parameters have a more accurate (i.e. most-likely prediction being closer to the true value) and more precise prediction (narrower distributions) with the inclusion of second-order moments. The distance functions are the same as those used in Figure 2.

The inference quality varies across different types. We examine the inference quality as a function of the type abundance (Figures S2 and S3). For the Lotka-Volterra model with 3-types and absolute abundances, the growth rates *f* and self-interaction terms *I*_*aa*_ tend to be more accurately inferred for the more abundant types. The cross-interaction terms *I*_*ab*_ are less accurately inferred overall, even when taking into account the first- and second-order moments (Figure S3). For the logistic model with 5-types and absolute abundances, the growth rates *f* tend to be more accurately inferred for the more abundant types, and the immigration rates *m* tend to be inferred to a similar degree of accuracy irrespective of the abundance of the type. The inference qualities for growth rates and immigration rates are improved with the incorporation of second-order moments, however the death rates *ϕ* in the logistic model are not accurately inferred (Figure S2).

### 3.2 Caveats associated with different distance functions and stopping criteria

The combination of first-order moments and second-order moments in the distance function is not straightforward, as they differ substantially in magnitude. Since second-order moments are the average of products of two first-order moments, their contribution to the distance function must be rescaled by a factor of half to enable a proper comparison across moment orders. Several rescaling approaches exist for the second-order moments, each yielding a different distance function with different sensitivities to second-order contributions. The distance functions used in this work are listed in Table 1. In Figure 4, we compare the inference quality and minimization trajectories obtained using different distance functions and different stopping criteria, for the Lotka-Volterra model with 3 types and absolute abundances (the corresponding figure with the logistic model is Figure S8). The resulting inference qualities with each distance function are shown in the left column of Figure 4 (panels A and C). The right columns of Figure 4 (panels B and D) shows the minimization trajectories of different distance functions, associated with a particular time-series. Different distance functions allow differing levels of minimization between the observed and predicted time-series, which directly influences parameter inference and the overall inference quality.

**Figure 4:**
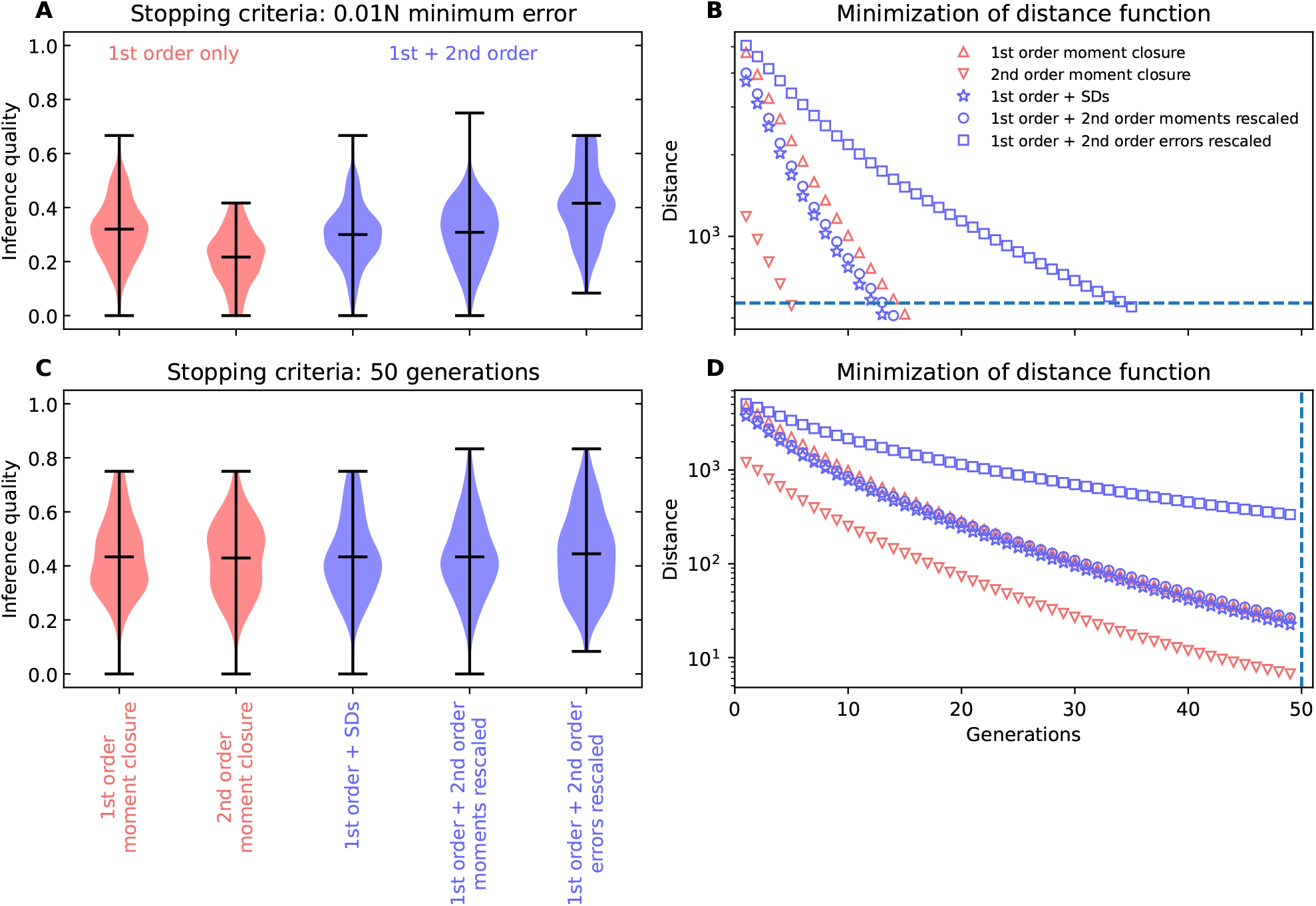
Comparison of inference qualities and minimization trajectories with different distance functions and stopping criteria for the Lotka-Volterra model with 3-types and absolute abundances. The top panels (A, B) show results with a stopping criterion of the distance function reaching a particular error threshold, while the bottom panels (C, D) utilize a stopping criterion of 50 generations in the ABC-SMC inference procedure. The left panels (A, C) depict inference qualities for different distance functions, while the right panels (B, D) shows minimization of different distance functions across generations associated with a particular time-series. Even though the extent of minimization varies across distance functions, especially with the stopping criterion used in panels (A, B), the inference quality is greater when the error terms of the second-order moments are rescaled before being combined with those from the first-order moments.

The top row of Figure 4 (panels A and B) corresponds to the case in which optimization is stopped when the distance function reaches a given threshold. We observe better inference quality results when the error terms of the second-order moments are rescaled before being combined with those from the first-order moments. Rescaling the errors allows for a greater degree of minimization, as depicted by the blue squares in Figure 4B. In the bottom row of Figure 4 (C and D), optimization is stopped after a fixed number of 50 generations in the ABC-SMC inference procedure. Under this stopping criterion, most distance functions give rise to similar inference qualities. However, some distance functions exhibit a greater degree of minimization than others, which may not be achievable in datasets subject to additional sources of noise beyond demographic stochasticity.

It is not straightforward to find a natural way to combine information from the first-order moments and second-order moments, as the inference quality varies considerably based on both the choice of distance function and the stopping criteria used in the optimization procedure. Because differing degrees of minimization lead to different inference qualities, establishing a fair and consistent metric to compare distance functions remains an open question.

## 4 Discussion

In this work, we have focused on a stochastic formulation for the dynamics of a microbial community, which describes how the abundances of different microbes change over time. We inferred the parameters of the underlying ecological model by fitting the first- and second-order moments of the observed time-series to those predicted by the model. We have shown under which conditions a larger fraction of the parameters are correctly inferred by extracting additional information from the dynamics of second-order moments.

For absolute abundances, we have shown that including second-order moments increases inference quality for both the logistic and Lotka-Volterra models (Figure 2). These results depend on the chosen distance function and the stopping criteria (Figure 4 and Figure S8). Additionally, the inclusion of second-order moments further collapses the posterior distributions (Figure 3 and Figure S6), indicating increased certainty for the inferred parameters. With the inclusion of more types, the fraction of correctly inferred parameters decreases. It should be noted that, while the number of parameters to be inferred increases with the number of types (linearly in the case of the logistic model, and quadratically in the case of the Lotka-Volterra model), we did not proportionally increase the number of time points, in order to be consistent with the typical temporal resolution of empirical microbiome datasets. And even with longer time-series this problem may not necessarily be resolved. Numerous studies have addressed this issue using various approaches: For example, Ref. [11] provides the option of reducing the number of types by grouping them together into ‘interaction modules’ based on similar behavior. Ref. [8] treats all the less-abundant types besides the top 10 most-abundant as ‘other’. However, Ref. [6] has cautioned against this kind of dimensionality reduction, showing that removal as well as grouping of less-abundant types leads to poor inference results on a synthetic dataset. Along similar lines, we find that parameters associated with less-abundant types tend to be less accurately inferred (Figure S2 and S3). This suggests that for datasets with a large number of types, it is harder to accurately infer the ecological processes and interactions involving the less-abundant types.

Working with relative abundances is more challenging than absolute abundances. As shown in Figure S5, the initial population size *n*_0_, a key parameter, is not inferable at all. However, advances in quantitative microbiome profiling (via flow cytometry or qPCR [26]) allows us to estimate *n*_0_, thereby enabling a transformation from relative to absolute abundances even as populations continue to vary, as described in Appendix B.

More broadly, some models may not be globally identifiable, i.e. multiple model parameter sets may give rise to identical time-series [6], which implies that accurate time-series prediction does not beget accurate parameter inference. This issue may be rectified by various means, including parameter transformations, using a stochastic model, log-transforming or refining prior distributions, removing weakly identifiable parameters by estimating them via alternative approaches, or switching to a globally identifiable model. Computational tools such as GenSSI [27] exist for determining the structural identifiability of a model, i.e. whether the model parameters are globally or locally identifiable given infinite, noiseless data. For example, among the two models we considered, the Lotka-Volterra model is globally identifiable whereas the logistic model is only locally identifiable [9]. Although we have not explored this aspect here, Bayesian methods offer built-in techniques for model selection, identifying the most probable and parsimonious model. Hence, parsimony and identifiability can be combined to establish a holistic framework for model selection and parameter inference.

While fitting models to data and inferring parameters, it is worth considering how accurately these models capture the underlying ecological processes. Ref. [28] argue that the Lotka-Volterra model is an over-parametrized description of the system, and cautions against its use for parameter inference from data. On the other hand, the Lotka-Volterra model, in its generalized form, only considers pairwise interactions, and extended versions incorporating additional complexities and higher-order interactions have been proposed [29–33]. While these approaches may capture the underlying processes and interactions to a greater extent, this comes with the added cost of inferring additional parameters. At the same time, the optimization procedure may utilize these additional degrees of freedom to overfit to the data. Therefore, it is challenging to find a balance between these competing narratives.

While we focus on using stochastic models and demographic noise as a conduit to extract more information from the system, we are ignoring other sources of variability, such as environmental noise and measurement noise. Typically, microbiome time-series data feature a lot of variability across replicates, which are likely due to a combination of environmental noise and limitations of the measurement technique (if a type goes extinct, its population has probably just decreased below the detection limit).

In such cases, it can become necessary to move beyond time-series-based distance measures and instead fit to appropriate summary statistics. In the same spirit, Ref. [28] advocates for greater focus on aggregate state variables, such as diversity indices. However, even the choice of the diversity index is not as straightforward as it may seem, as different indices capture different aspects of the system [34, 35]. Using one particular diversity index, e.g. Shannon diversity, as a distance measure discards a lot of information in a system that is already inherently complex. This is not a good choice given that ABC is known to be perform poorly with less-informative summary statistics [36]. For a comprehensive review of statistical methods for microbiome time-series analysis, see Ref. [37].

As far as the method of parameter inference is concerned, Approximate Bayesian Computation is often the first approach utilized, especially when the likelihood does not exist in analytical form (e.g. stochastic models, coalescent trees, etc). Finally, future work should in addition incorporate environmental noise while simultaneously working with the large number of microbial types typically found in microbiomes.

## 5 Acknowledgements

We thank the Department of Theoretical Biology in Plön for their numerous insightful suggestions. We also thank Román Zapién-Campos and Florence Bansept for their comments on our work.

## 8 Author Contributions

- **Conceptualization:** Chandan Relekar, Amanda de Azevedo-Lopes, Michael Sieber, Arne Traulsen.
- **Data curation:** Chandan Relekar, Amanda de Azevedo-Lopes.
- **Formal analysis:** Chandan Relekar, Amanda de Azevedo-Lopes.
- **Investigation:** Chandan Relekar, Amanda de Azevedo-Lopes, Michael Sieber, Arne Traulsen.
- **Methodology:** Chandan Relekar, Amanda de Azevedo-Lopes, Michael Sieber, Arne Traulsen.
- **Software:** Chandan Relekar
- **Supervision:** Arne Traulsen
- **Validation:** Chandan Relekar, Amanda de Azevedo-Lopes, Michael Sieber, Arne Traulsen.
- **Visualization:** Chandan Relekar, Amanda de Azevedo-Lopes, Michael Sieber, Arne Traulsen.
- **Writing - original draft:** Chandan Relekar
- **Writing - review and editing:** Chandan Relekar, Amanda de Azevedo-Lopes, Michael Sieber, Arne Traulsen.

## 9 Data Availability

The code and data can be found at https://doi.org/10.5281/zenodo.22898948

## A Expressions for first-order moments and second-order moments

The expressions below were first derived in Ref. [9]. The systems of ODEs were simulated using the solve ivp function of SciPy. LSODA was used as the method of integration for both models. Stoppers were utilized in case the populations went negative, or started to diverge. For the first-order moments, only the dynamical equations for the first-order moments were used (Eqs. S1 and S4), and the second-order terms present in them were approximated (e.g. ⟨*n*_*k*_*n*_*l*_⟩ ≈ ⟨*n*_*k*_⟩ ⟨*n*_*l*_⟩). With first- and second-order moments, all the equations below were used, and the third-order terms were approximated in the dynamical equations of the second-order moments (e.g. 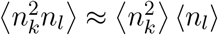).

### A.1 Logistic model

By plugging in the growth and death rates from Eqs. 1 and 2 into the Master equation (Eq. 3) and using Eq. 4 to derive the equations for the statistical moments (by setting 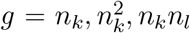), we obtain the expressions for the first-order moments (means) and second-order moments (square- and co-) to be

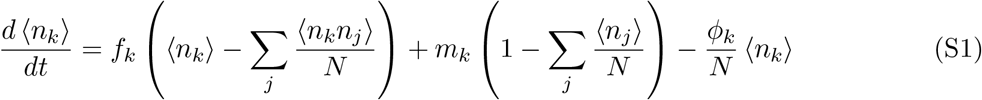

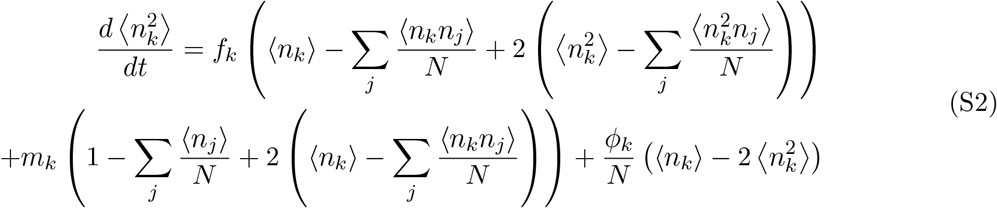

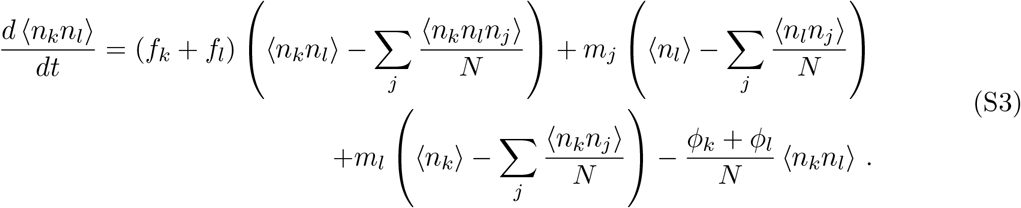

### A.2 Lotka-Volterra model

Repeating the same procedure as before with Eqs. 6 and 7, we obtain

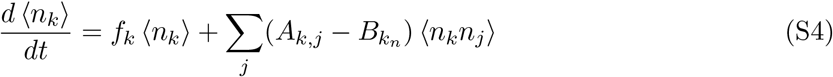

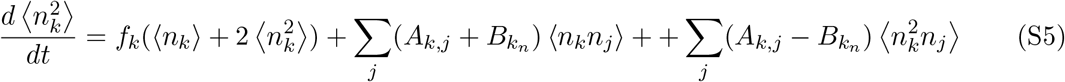

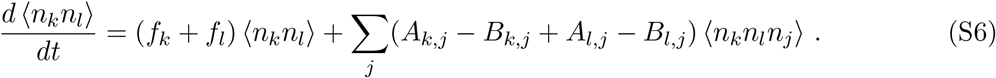

## B Working with relative abundances

Often empirical measurements of microbiome composition are given in terms of relative abundances and not as absolute abundances. Our framework also allows us to rewrite our equations to work with relative abundances. In this case, the initial population *n*_0_ becomes an additional model parameter that either has to be inferred or provided. Let 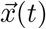 be used to denote relative abundances, and *n*_Σ_(*t*) denote the total population at time *t*. Thus, we can write the absolute abundances as

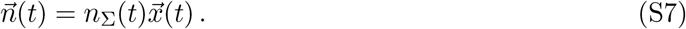

On differentiating the above equation and using chain rule, we obtain derivatives for the moments in terms of relative abundances, which come out to be [9]

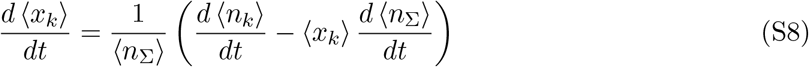

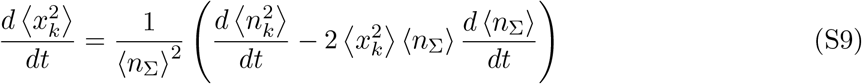

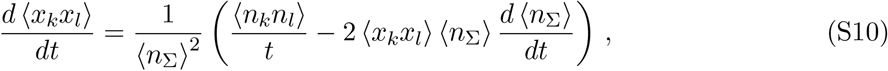

where

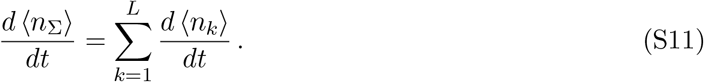

Starting from relative abundances at the first time-point,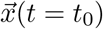, we apply the transformation in Eq. S7 using *n*_Σ_(*t* = *t*_0_) = *n*_0_ in order to obtain absolute abundances,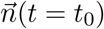. Using the equations in Appendix A.1 and A.2, we estimate the absolute abundances at the next time step (*t*_1_) and sum them up to obtain *n*_Σ_(*t* = *t*_1_). Finally, we convert the derivatives of absolute abundances to those in relative abundances using Eqs. S8, S9 and S10, which we subsequently use to estimate relative abundances at the next time-step. These steps are repeated at the subsequent iterations. Figure S1 presents results analogous to those in Figure 2, but for time-series data expressed as relative abundances. It illustrates the greater difficulty in accurately inferring model parameters in this setting. Figure S5 indicates that the initial population, *n*_0_, is structurally unidentifiable when working with relative abundances. Hence, it is preferable to measure *n*_0_ during the experiment. Notably, no clear trends are observed with respect to either the inclusion of higher-order moments or number of types, in either model.

## C Priors

All prior distributions are uniform. Gaussian priors were avoided to ensure compatibility with potential log-transformations of the parameter space. Moreover, uniform priors correspond to the case where, beyond the range of biological plausibility, we have no prior information of the distribution a parameter should follow. Hence, uniform priors provide a natural and well-defined baseline for Bayesian inference. The prior ranges for each parameter are listed in Tables S1 and S2, and were chosen to ensure that the resulting time-series exhibit biologically meaningful and dynamically rich behavior. The initial population size *n*_0_ is included as a parameter only when working with relative abundances.

## D Hyper-parameters for ABC

### Population

The population of particles used during parameter inference is set to 1000 *×L*, where *L* is the number of types.

### Transition Kernel

For the logistic model, the Multivariate Normal Transition (MNT) kernel was used with default parameters. For the Lotka-Volterra model, the Local Transition kernel was used with default parameters.

### Minimum acceptance threshold *ϵ*

The minimum acceptance threshold *ϵ* was calculated to ensure consistent and comparable stopping conditions across all 100 inference runs using randomized datasets and across all number of types, *L*. The values below were chosen based on the trade-off between accuracy of inferred parameters and feasibility of inference

- For the logistic model with absolute abundances, *ϵ* = 0.01*N* was used, where *N* is the carrying capacity associated with the corresponding dataset.
- For the Lotka-Volterra model with absolute abundances, *ϵ* = 0.03 *×* max(*{n*_*avg*_(*t*)*}*), where 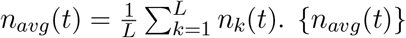 is the set across all *t* = *t*_0_, *t*_1_, …, *t*_*m−*1_.
- For both models with relative abundances, *ϵ* = 0.01 was used.

### Simulation details

We ran 100 inference runs corresponding to 100 randomized datasets for each number of types *L*. Sufficient RAM was allocated for each run (25 GB on a cluster with 64 nodes), alongside a time limit of two days. For the number of types *L* that we worked with, it was exceedingly rare for a job to exceed these constraints. PyABC version 0.12 was utilized for inference.

## E Generation of randomized time-series

To assess trends in identifiability, we fitted multiple synthetic datasets and computed the average observed accuracy of parameter inference across them. For each number of types *L*, we randomly sampled 100 model parameter sets from the prior distributions. We used only those time-series which exhibited no unbounded growth (defined as absolute abundance of any type exceeding 10^16^), no extinctions (defined as the absolute abundance of any type falling to zero or below), and relative abundances lying within biologically plausible bounds (0 ≤ *x*_*k*_ ≤ 1).

For each parameter set, we generated four independent replicates of synthetic time-series data using the Gillespie algorithm [19], with transition rates defined in Eqs. (1) and (2) for the logistic model, and Eqs. (6) and (7) for the Lotka-Volterra model. Each time-series was sampled at ten evenly spaced time points, as empirical datasets often contain around the order of ten time points. The observed first and second-order moments were computed by averaging over the four replicates. For the logistic model, this process was repeated for *L* = 2, 3, …, 7 types, and for the Lotka-Volterra model, this process was repeated for *L* = 2, 3, 4 types.

## F Additional figures

**Figure S1:**
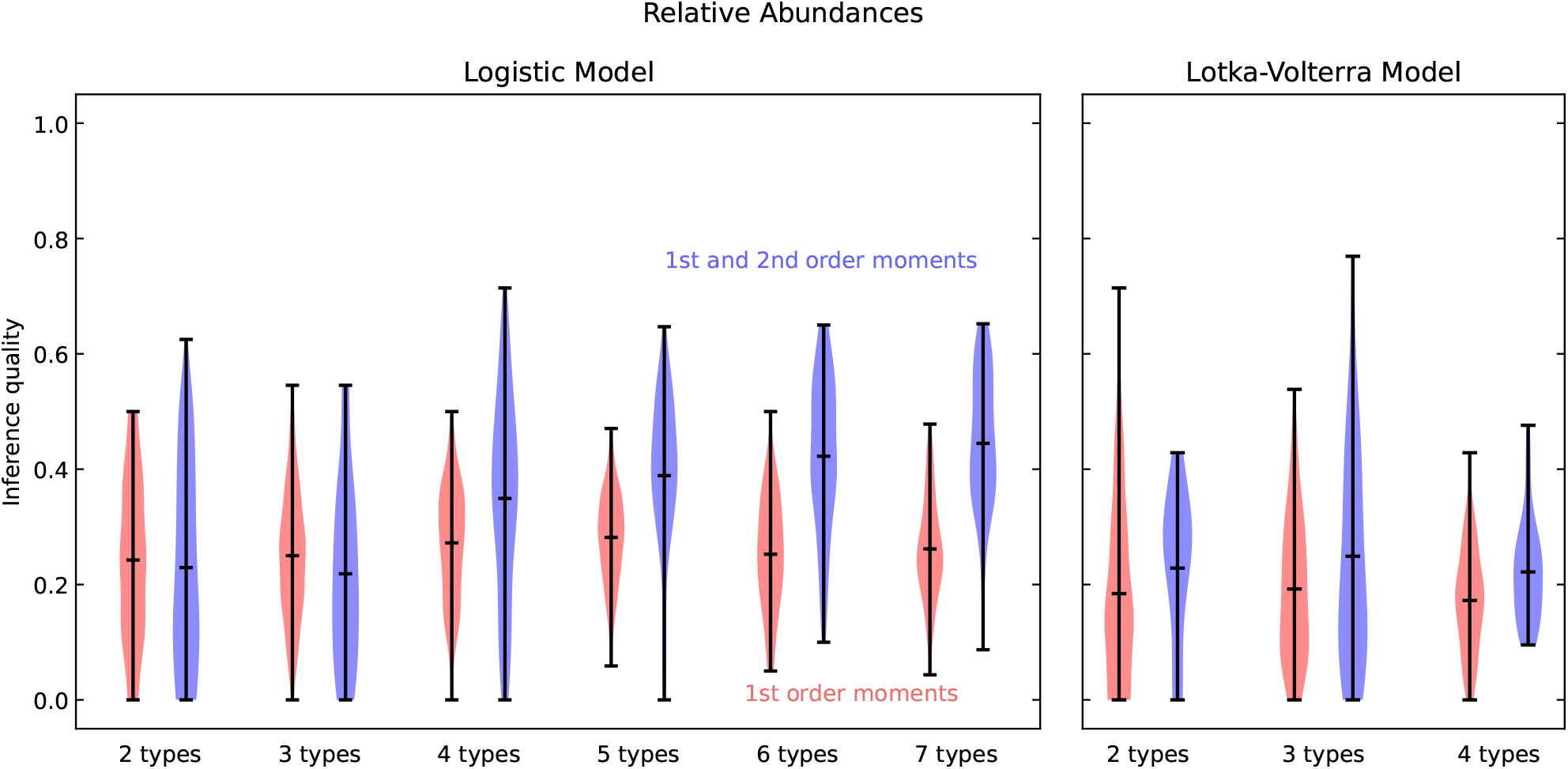
Estimating the inference quality on synthetic data with relative abundances. We compare the inference quality of fitting just first-order moments to the inference quality of fitting both first- and second-order moments, across 100 randomized datasets, for the logistic and the Lotka-Volterra models and different numbers of types, *L*. We do not observe any increase in inference quality due to incorporation of second-order moments. The distance functions are the same as those used in Figure 2.

**Figure S2:**
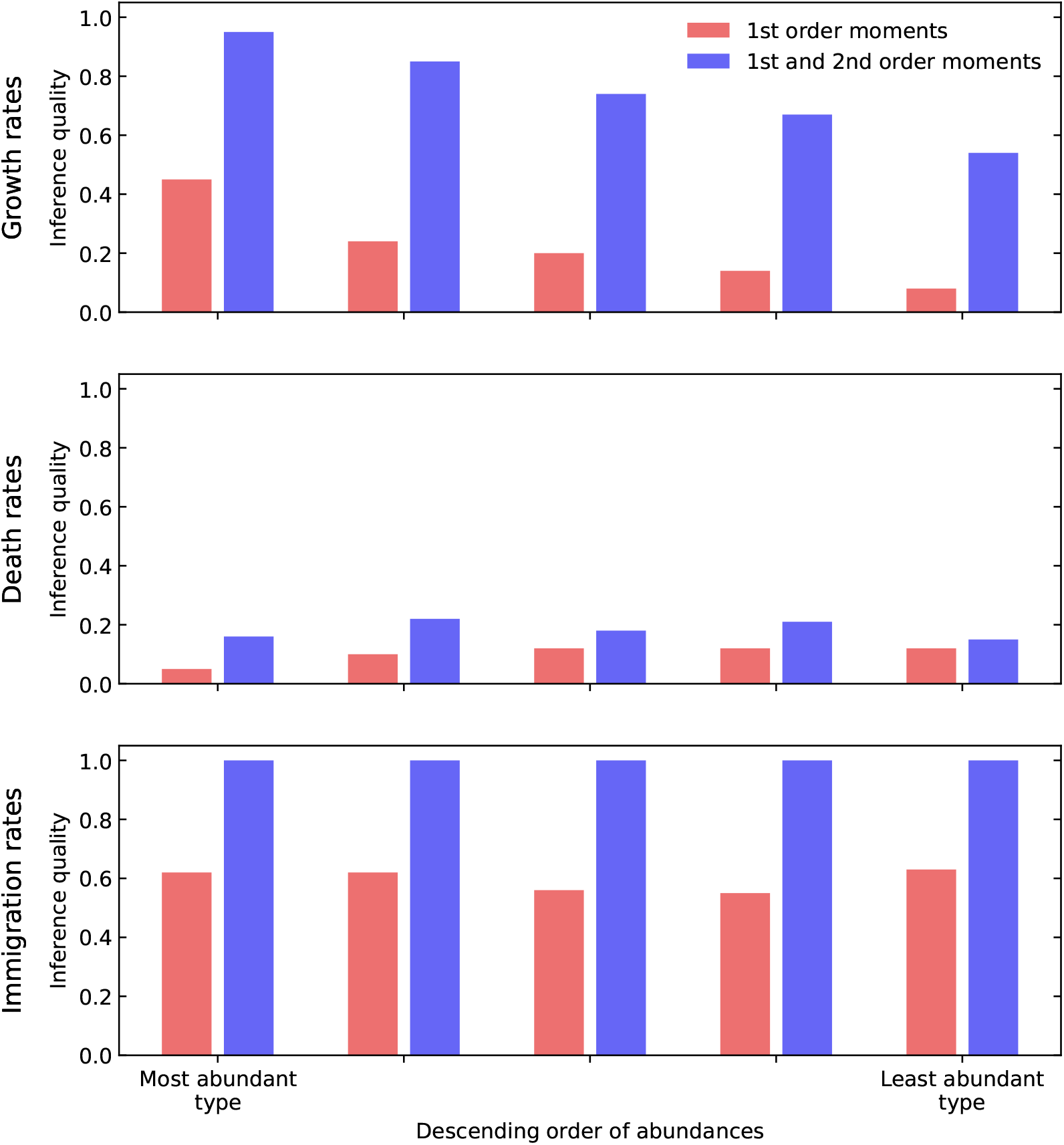
Comparing parameter-wise inference quality for decreasingly abundant types in the microbiome, with and without the inclusion of second-order moments. The different types are sorted according to their abundance at the last time-point. This figure is a further breakdown of the case of five types in the logistic model from Figure 2. Growth rates are more accurately inferred for the types that are more abundant.

**Figure S3:**
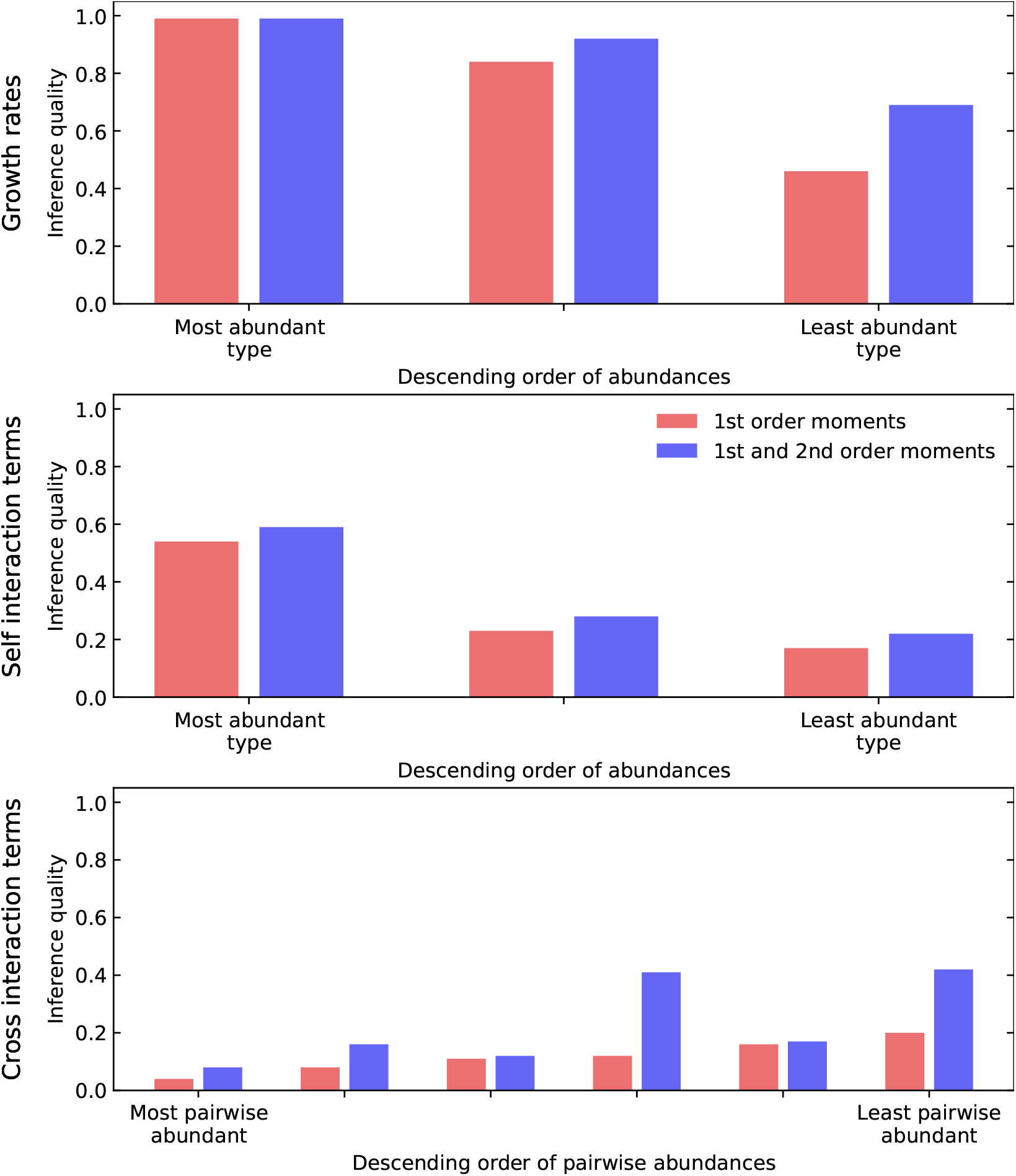
Comparing parameter-wise inference quality for decreasingly abundant types in the microbiome, with and without the inclusion of second-order moments. This figure is a further breakdown of the case of three types in the Lotka-Volterra model from Figure 2. The different types are sorted according to their time-averaged abundance. For sorting cross-interactions, we looked at the product of time-averaged populations of the involved types, and slightly biased the focal type. Growth rates and self-interaction terms are more accurately inferred for the types that are more abundant.

**Figure S4:**
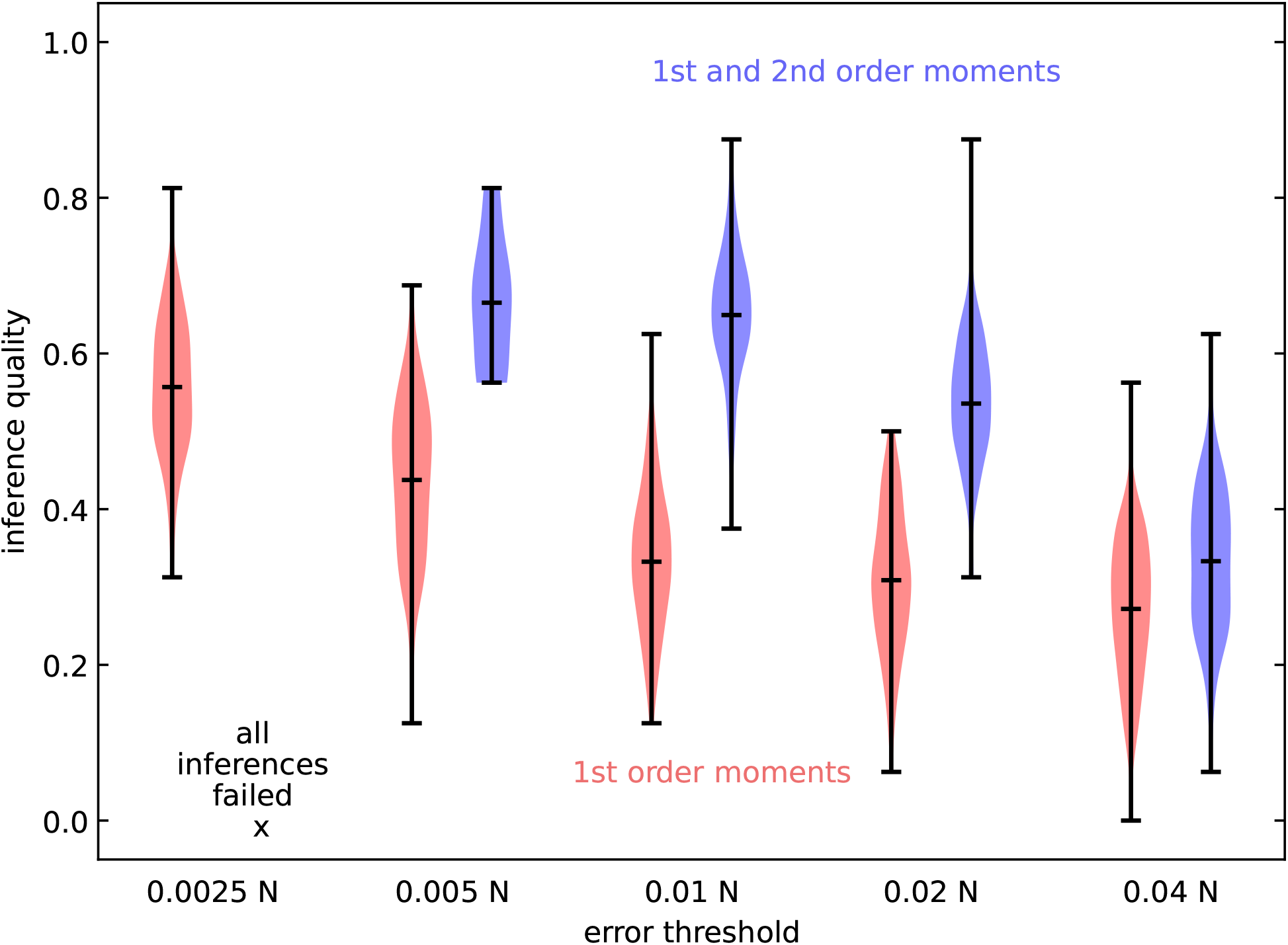
Comparing the inference quality for different error thresholds. We compare the inference quality for different error thresholds on synthetic data for the logistic model with five types and absolute abundances. As expected, the inference quality increases with decreasing error threshold for both first-order moments and first- and second-order moments. However, for the lowest error threshold, no inference run was able to reach the threshold with first- and second-order moments, and hence the inference quality is not reported for this case. The inference quality obtained by fitting only first-order moments to a very low error threshold (e.g., the red violin for *ϵ* = 0.0025*N*) is comparable to the inference quality achieved when fitting first- and second-order moments to a higher error threshold (the blue violin for *ϵ* = 0.02*N*). However, fitting to such low error thresholds (e.g. *ϵ* = 0.0025*N*) is only feasible with synthetically generated datasets, whereas empirical datasets not only contain additional sources of noise but may exhibit substantial deviation from theoretical models. The distance functions are the same as those used in Figure 2.

**Figure S5:**
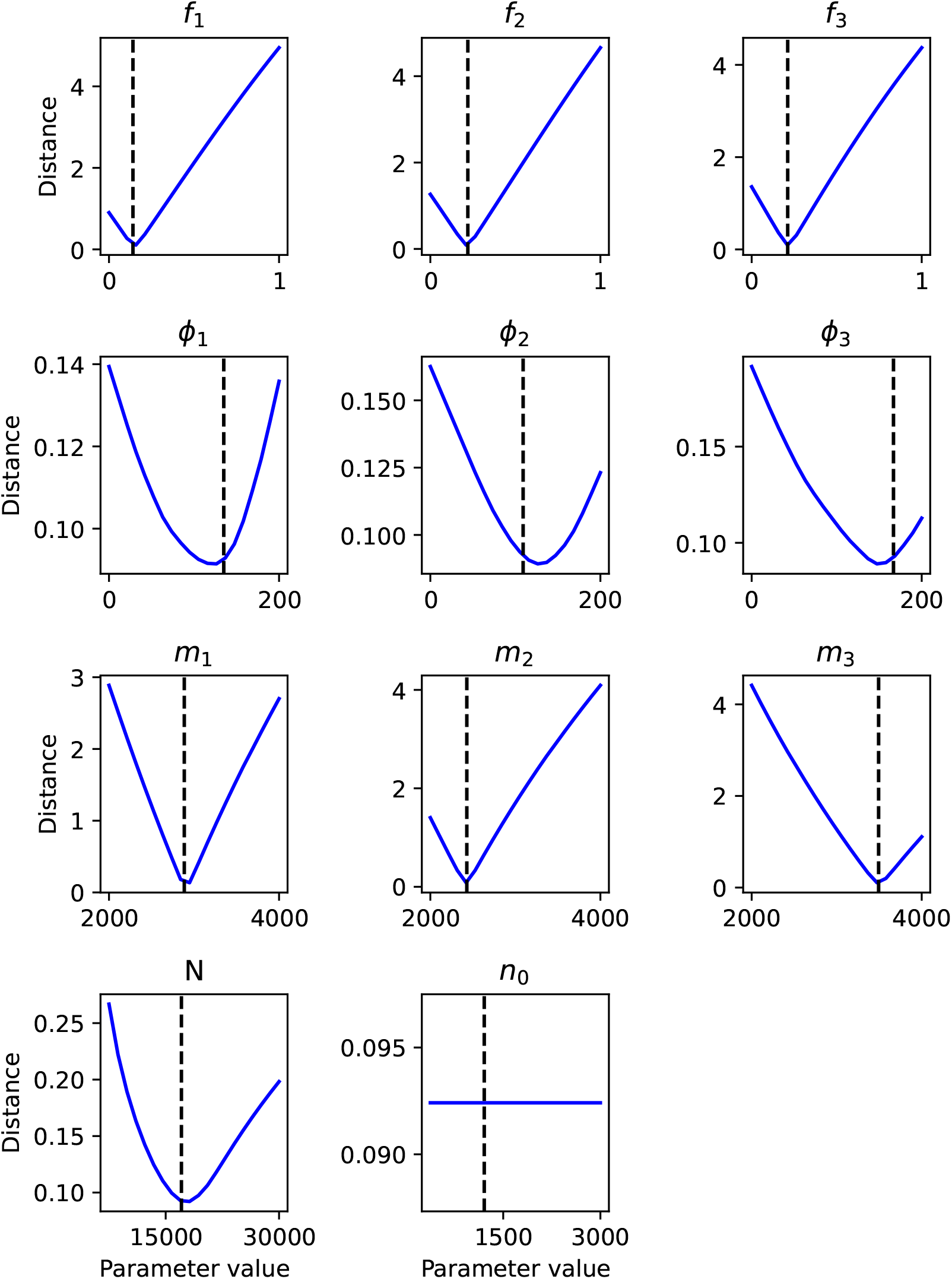
Variation in distance function of the logistic model with three types and relative abundances, as each individual parameter is varied while other parameters are set to their true values. This suggests that the death rates *ϕ* are less identifiable, and the initial population *n*_0_ is not at all identifiable. The ‘first-order and second-order errors rescaled’ distance function is used.

**Figure S6:**
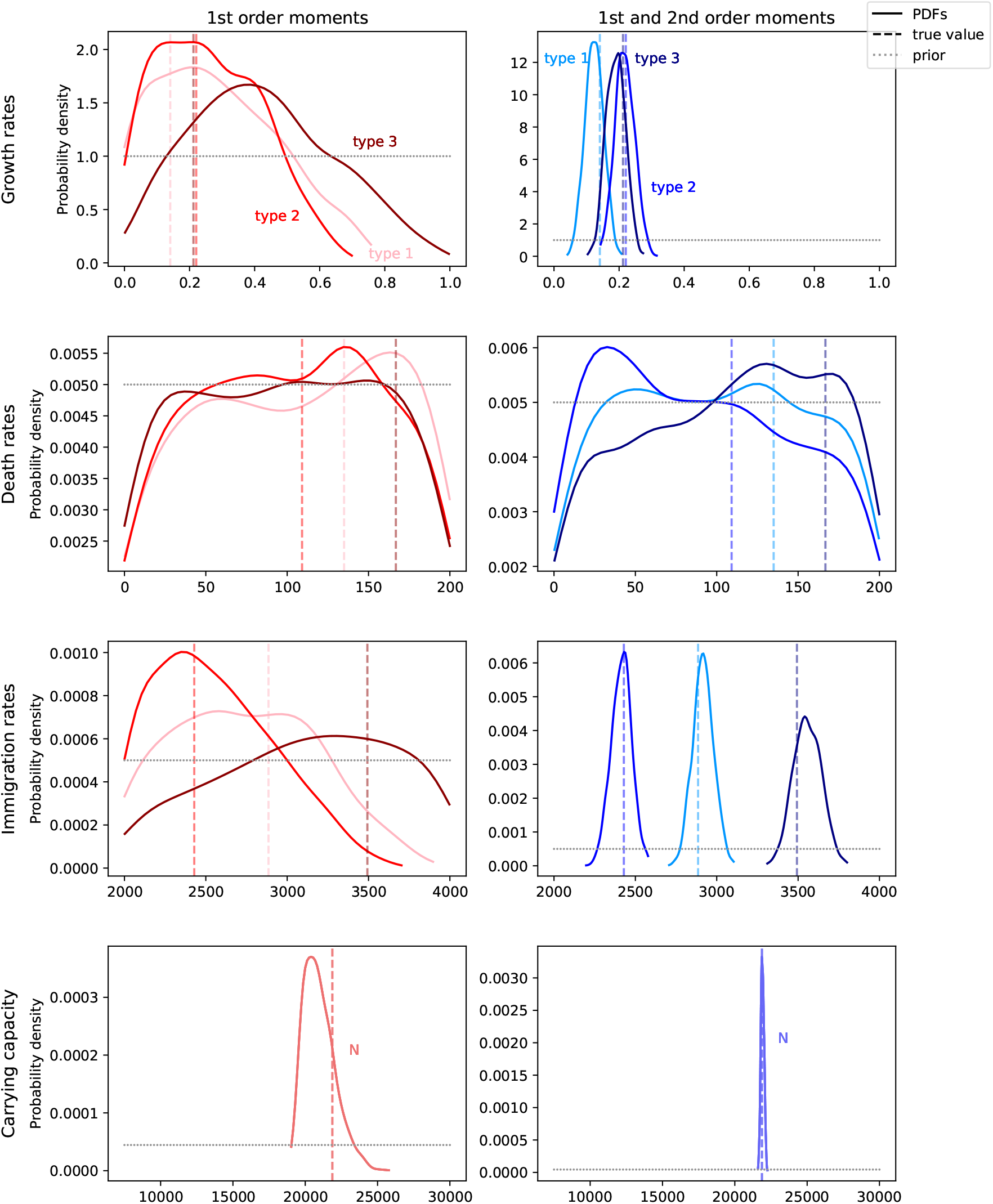
Comparison of the posterior distributions with and without the inclusion of second-order moments for the logistic model with three types and absolute abundances. Except for the death rates, all parameters have a more accurate (most likely prediction being closer to the true value) and more precise prediction (narrower distributions) with the inclusion of second-order moments. The distance functions are the same as those used in Figure 2.

**Figure S7:**
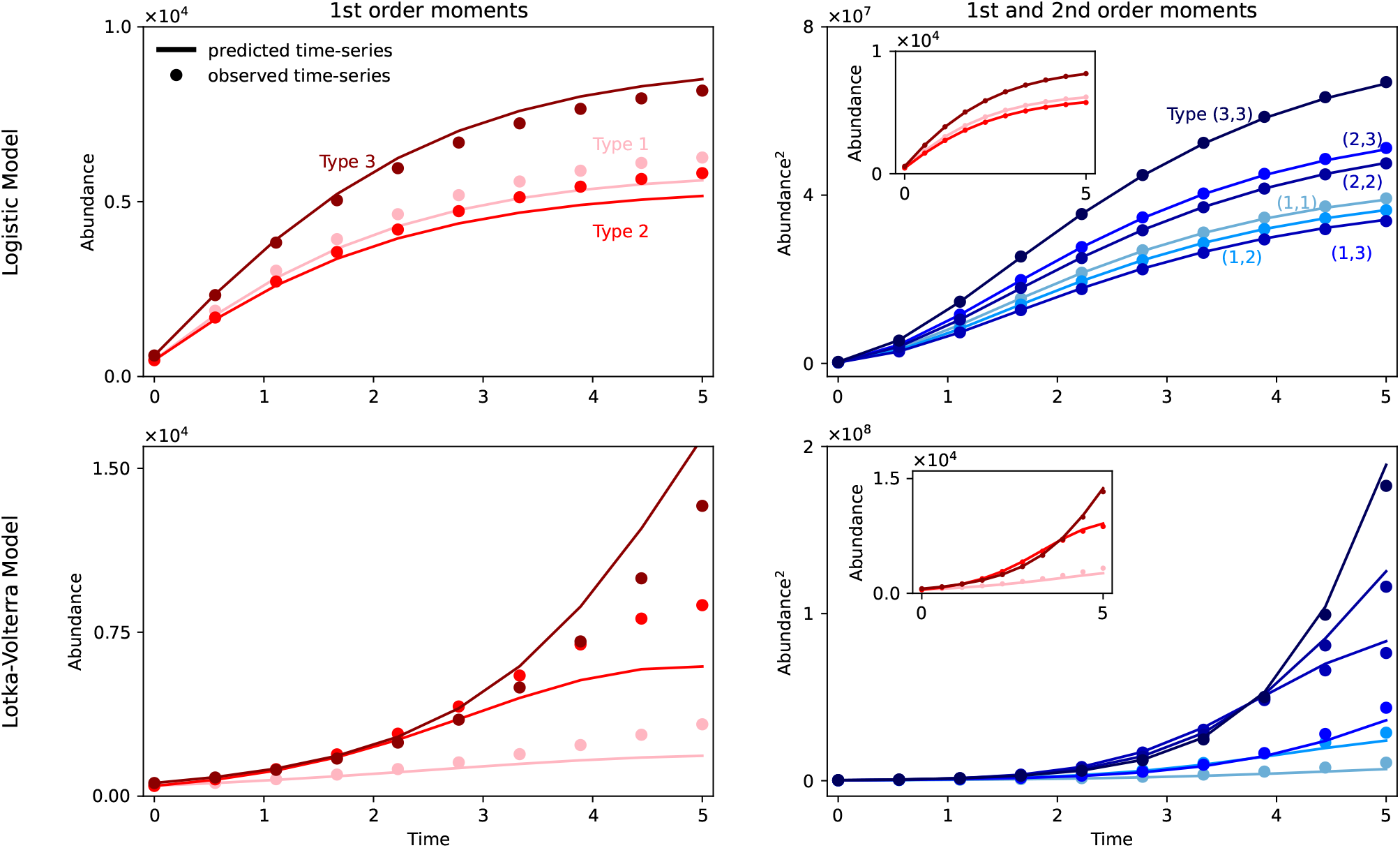
Original time-series (points) vs. predicted time-series (lines) resulting from the most-probable model parameters from Figure 3 (depicted in the top row of the above figure) and Figure S6 (bottom row). The left column showcases the result whilst utilizing just the first-order moments. The right column depicts the result with the additional inclusion of second-order moments too (the corresponding first-order moments are shown in the inset).

**Figure S8:**
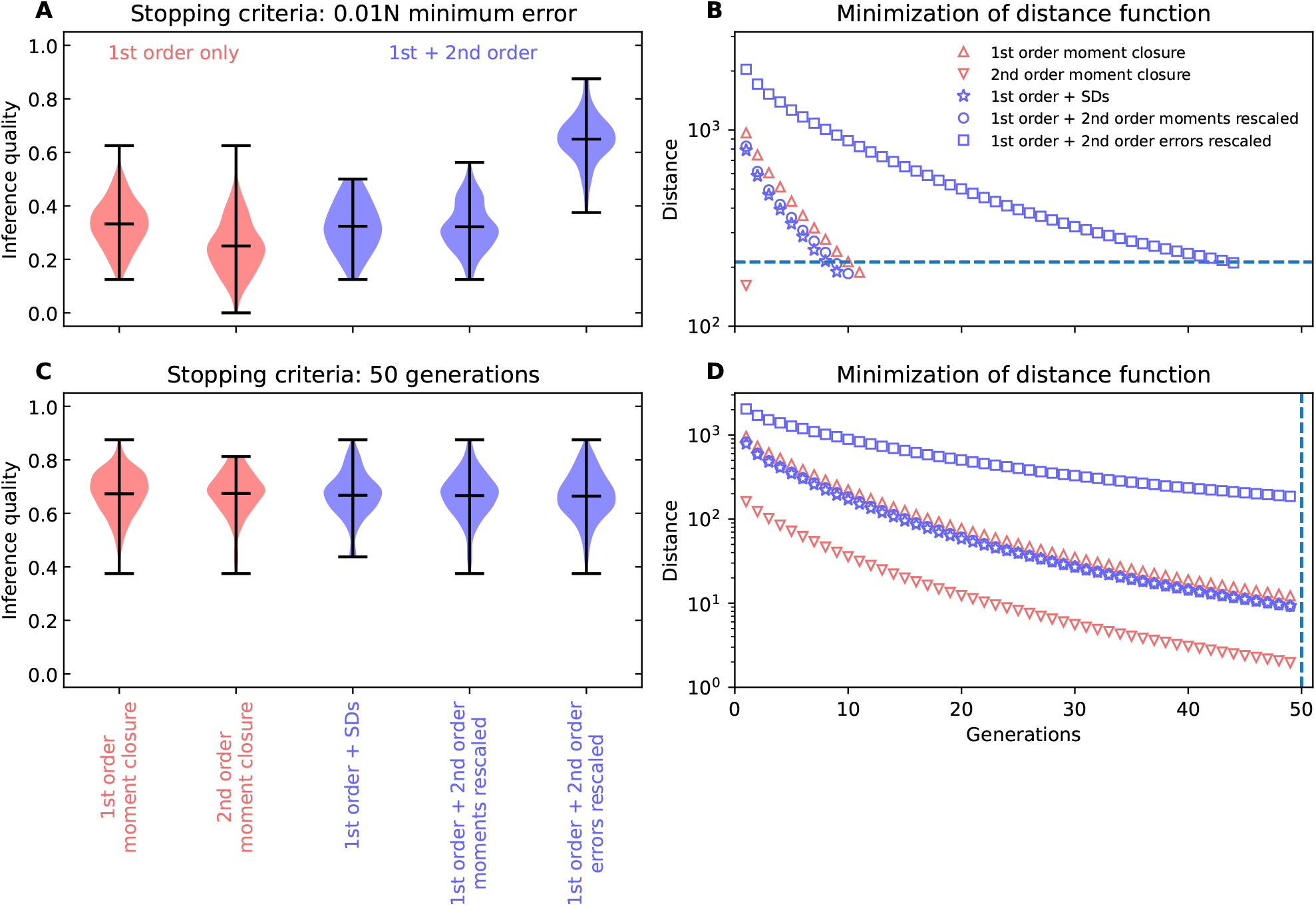
Comparing inference qualities and minimization trajectories resulting from using different distance functions and different stopping criteria. The top panels (A, B) show results with a stopping criterion of the distance function reaching a particular error threshold, while the bottom panels (C, D) utilize a stopping criterion of 50 generations in the ABC-SMC inference procedure. The left panels (A, C) depict inference qualities for different distance functions, while the right panels (B, D) shows minimization of different distance functions across generations associated with a particular time-series.

**Table S1:** Priors for the logistic model used during Bayesian inference. These priors were also used to generate synthetic datasets for the logistic model.

| Parameter | Range | Units |
| --- | --- | --- |
| Growth rates $f$ | $(0, 1)$ | $\text{time}^{-1}$ |
| Death rates $\phi$ | $(0, 200)$ | cells / time |
| Immigration rates $m$ | $(2000, 4000)$ | cells / time |
| Initial population $n_0$ | $(125L, 1000L)$ | cells |
| Carrying capacity $N$ | $(2500L, 10000L)$ | cells |

**Table S2:** Priors for the Lotka-Volterra model used during Bayesian inference. These priors were also used to generate synthetic datasets for the Lotka-Volterra model.

| Parameter | Range | Units |
| --- | --- | --- |
| Growth rates $f$ | $(0, 1.5)$ | $\text{time}^{-1}$ |
| Self-interaction $I_{aa}$ | $(-10^{-4}, 0)$ | $(\text{cells} \cdot \text{time})^{-1}$ |
| Cross-interaction $I_{ab}$ | $(-5 * 10^{-5}, +5 * 10^{-5})$ | $(\text{cells} \cdot \text{time})^{-1}$ |
| Initial population $n_0$ | $(125L, 1000L)$ | cells |

